# Cytokinin acts as a systemic signal coordinating reproductive effort in Arabidopsis

**DOI:** 10.1101/2024.07.09.602718

**Authors:** Darya Volkava, Sona Valuchova Bukovcakova, Pavlina Mikulkova, Jana Pecinkova, Jan Skalak, Jan Hejatko, Karel Riha

## Abstract

The duration of the flowering phase is a key adaptive trait related to reproductive effort and plant longevity. In plants with indeterminate inflorescences, the number of flowers produced is determined by the activity of the inflorescence meristem. However, the integration of inflorescence meristem activity with the physiological and developmental status of a plant is not well understood. We combined quantitative imaging of the inflorescence meristem with the manipulation of cytokinin levels and distribution to investigate how meristem activity is regulated. Our data support the hypothesis that emerging fruits drain the acropetal flow of cytokinin away from the meristem inducing its proliferative arrest. We further show that the duration of flowering and cytokinin signaling in the inflorescence meristem is influenced by nitrate availability. Taken together, these data suggest that cytokinin functions as a systemic signal that integrates information about ongoing reproductive effort and plant nutrient status to control flower production.

## Introduction

In contrast to animals, plants exhibit a remarkable flexibility in their body structure, continuously generating new organs and tissues through post-embryonic growth. This developmental plasticity allows plants to dynamically adapt their physical structure to local environmental conditions and adjust their life strategy accordingly (1). Central to this process is flower production, a key developmental trait that is directly related to reproductive effort. The timing and duration of the flowering phase determine seed yield, quality, and reproductive success. However, allocating resources to fruit and seed production is an energetically demanding task, so deciding how many flowers to produce must be integrated with developmental and environmental cues (2, 3).

In plants with indeterminate inflorescences, the number of flowers is limited by proliferative arrest (PA) of the inflorescence meristem, which occurs after production of a certain number of fruits. A major factor controlling the arrest is seed production, since surgical removal of fruits or mutations causing infertility greatly extend the activity of the inflorescence meristem and the life span of the plant (4–7). Two hypotheses have been proposed to explain the corelative control of flower formation by developing seeds (8–10). The “death hormone” hypothesis predicts the existence of a factor produced by the fruit or seed that is transported to other tissues where it induces the cessation of growth. The “nutrient drain” hypothesis suggests that developing reproductive organs represent a strong sink that diverts resources or signaling molecules that sustain growth away from vegetative tissues.

The corelative inhibition between reproductive organs of the same plant was described in a number of species, but deeper insights into this phenomenon have been obtained only recently by studies in *Arabidopsis thaliana* (10–12). The primary inflorescence in Arabidopsis becomes inactive after production of ∼30-50 flowers and arrests with a cluster of unopened floral buds (6, 13). The cessation of flower development occurs in two steps. The inflorescence meristem stops initiating new flower primordia 2-3 weeks after bolting, which is followed by the full arrest of floral bud development several days later (14, 15). Transcriptomic analysis revealed that the arrested meristem and floral buds enter a dormancy-like state that can be reverted into an active state by fruit removal in a process that resembles release of axillary buds from apical dominance (16).

Several molecular pathways have been implicated in the inflorescence arrest. As the plant ages, the inflorescence meristem gradually diminishes in size, which is accompanied by the downregulation of the stem cell maintenance factor WUSCHEL (WUS) (14, 16–19). The age-dependent decline of WUS involves FRUITFUL/APETALA2 regulatory module (18, 20), but its coordination with reproductive processes remains unknown. Jasmonic and abscisic acid signaling also contributes to establishment of proliferative arrest and disruption of these signaling pathways extends the flower production (21, 22). The onset of floral arrest is highly susceptible to fruit formation in the vicinity of the inflorescence meristem (23). It has been proposed that the fruits proximal to the inflorescent meristem exert an inhibitory effect via auxin canalization. According to this model, increasing auxin flow from emerging fruits disrupts the polar auxin transport from inflorescence apex causing its arrest (14, 23).

However, the inhibitory effect of fruits on inflorescence apex becomes effective only when the meristem reaches a certain age and develops competence for arrest (23). This indicates existence of an additional mechanism that tracks the reproductive and developmental age of the meristem. Recent studies have shown downregulation of cytokinin signaling during inflorescence arrest, and that exogenous application of cytokinin to the inflorescence meristem delays or even reverts the arrest (15, 19). Furthermore, mutations in genes involved in cytokinin signaling significantly affect inflorescence arrest (15, 20, 24–27). In shoot apical meristem, cytokinin reinforces expression of WUS in a positive feedback loop and helps to maintain the undifferentiated state of stem cells (28). In addition, cytokinin stimulates cell proliferation and meristematic activity by promoting CYCLIN D expression (29). Thus, it is likely that cytokinin signaling plays a central role in regulating inflorescence arrest.

Cytokinins act as long-distance signaling molecules and some of their forms are produced in roots and transported via xylem into shoot to regulate growth and developmental processes (30, 31). Developing seeds and embryos exhibit high level of cytokinin, and at least part of it may be provided from maternal plant (32). Thus, the emerging fruits with developing seeds may represent a sink that diverts cytokinin transport from the inflorescence apex, leading to decline in cytokinin signaling in the inflorescence meristem and increasing its competence to proliferative arrest. In this study we used quantitative imaging to investigate whether cytokinin acts as a systemic signal that integrates physiological and developmental cues to coordinate flower production and reproductive efforts.

## Results

### Meristems become permissible to inflorescence arrest at older age

Under our experimental conditions, *A. thaliana* Col-0 produces in average 53 flowers on the primary inflorescence (PI) in about 25 days before proliferative arrest occurs (Supplementary Fig. 1). Continuous pruning of flowers on the PI substantially increased both the number of flowers and the duration of the flower production. The first 30 flowers have a negligible effect on the onset of inflorescence arrest and their removal did not significantly alter the duration of flowering or the number of flowers produced. However, continuous pruning of flowers appearing after the development of the first 30 flowers had the same impact as removing all flowers (Supplementary Fig. 1). These observations confirm the data of Ware et al. (23) and suggest that the inflorescence arrest is primarily influenced by the production of apex-proximal fruits and that it occurs only when the inflorescence meristem has passed a certain developmental age and becomes susceptible to the arrest.

### Quantitative imaging of inflorescence meristem

The observation of diminished cytokinin signaling in the arrested meristem (15, 19) raised the question of whether the declining transport of cytokinin to the inflorescence apex due to sink in developing fruits determines the susceptibility of the meristem to arrest. Such scenario predicts that cytokinin signaling in the meristem gradually decreases with the growth of new fruits. To test this hypothesis, we developed a methodology for three-dimensional (3D) quantitative imaging of fluorescent reporters in inflorescence meristems (IM) using light-sheet fluorescence microscopy (LSFM). In a standard confocal microscopy, the apical meristem is scanned from the apex to the base, and scanning of a complete confocal stack in multiple channels can take tens of minutes (33, 34). During image acquisition, the entire volume of the meristem is illuminated by the excitation laser, which can partially quench the signal, leading to inaccuracies in the signal quantification, especially in the deeper layers of the tissue.

LSFM allows fast 3D imaging of relatively macroscopic objects by passing them through a thin laser sheet, which minimizes the photodamage. To accurately capture the 3D architecture and to minimize the effect of 3D scanning on signal intensity in different layers, we imaged the inflorescence meristem laterally along the axis of vertical symmetry from eight different angles (Fig. 1a). The resulting stacks were combined to build a comprehensive 3D model of the inflorescence meristem (Supplementary Fig. 2, Fig. 1b, Movies 1 and 2). The intensity of each voxel in the model represents the average of the data obtained in the eight scans. We used two different approaches for data analysis (Supplementary Fig. 2). For the *pCLV3::H2B-mCherry* marker that gives a strong and well defined nuclear signal, we performed spot-based segmentation and included the mean intensity of the signal from all detected nuclei in the meristem for the downstream analyses (Fig. 1c-f). This provided an estimate of the size of the stem cell niche and the intensity of the CLV3 expression.

**Fig. 1.**
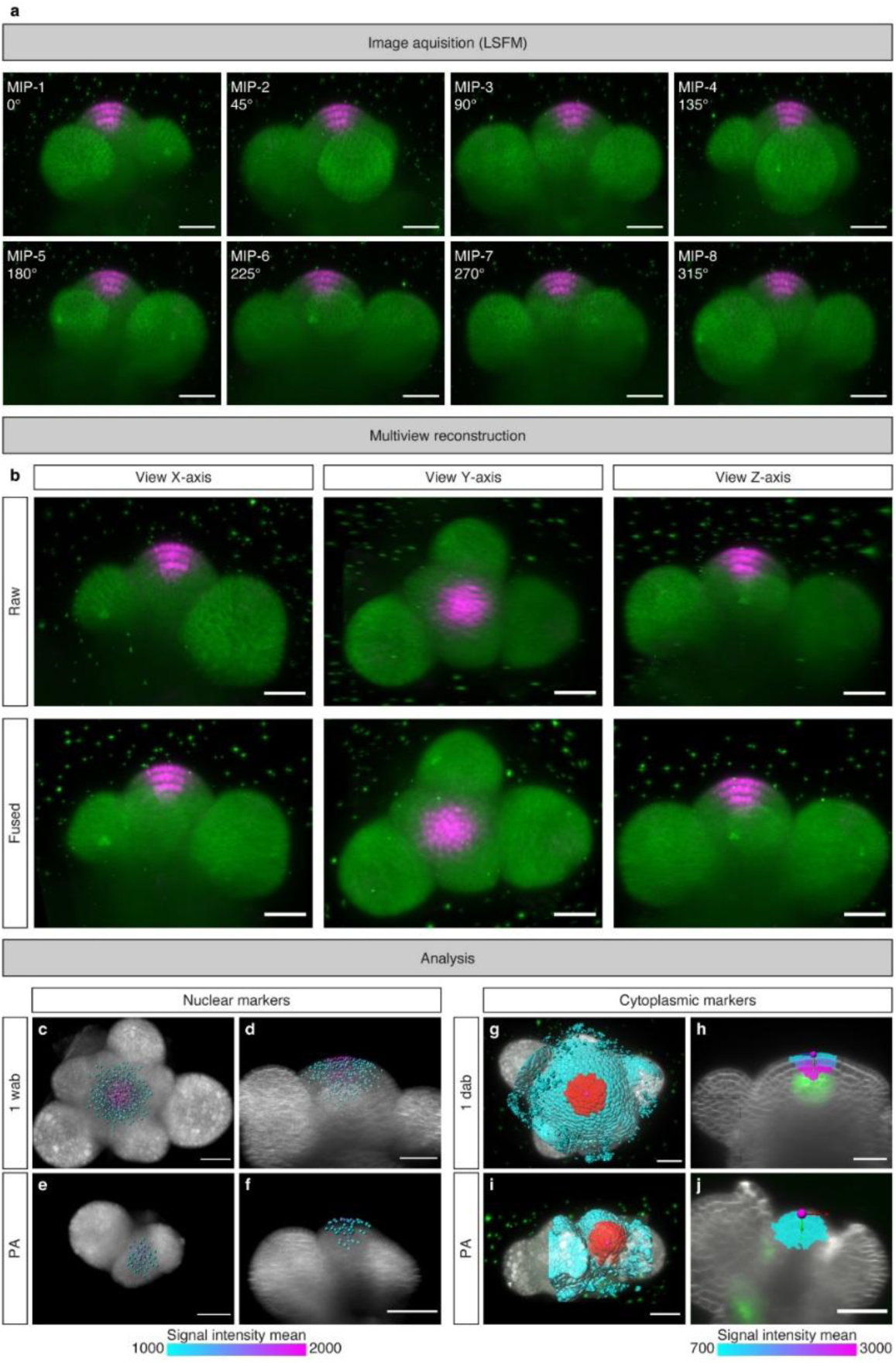
Quantitative 3D imaging of Arabidopsis inflorescence meristem. **(a)** Maximum intensity projections (MIPs) of IM expressing *pCLV3::H2B-mCherry* (magenta) and the plasma membrane marker *SYP132:GFP* (green) scanned from eight different angles. Scale bar = 30 µm. (**b)** Comparison of image quality from a single angle stack (above) and all angles fused together (below) **(c-f)** 3D-reconstructions of IMs 1 wab (c,d) and at PA (e,f) with the segmented nuclear *pCLV3::H2B-mCherry* signal. Relative signal intensity means of the segmented nuclei are indicated by colormap. Plasma membrane is highlighted by *SYP132:GFP* (gray). (c,e) represent top views and (d,f) longitudinal sections, respectively. Scale bar = 30 µm. **(g-j)** 3D-reconstructions of IMs 1 wab (g,h) and at PA (i,j) with cells segmented using plasma membrane signal obtained by FM4-64 staining (gray). (g,i), Top views of surface rendered models. Red color indicates cells within 21 µm from meristem apex that were used for quantitative analysis of cytoplasmic signal. (h,i) Longitudinal sections of the segmented IMs. Intensity means of the *TCSn::GFP* signal in cells within 21 µm from the apex are indicated by heatmaps. *TCSn::GFP* signal outside of that volume is displayed in green. A reference frame (magenta sphere with arrows) indicates position of the meristem apex. Scale bar = 30 µm.

The nuclear signal from *pWUS::WUS-linker-GFP* marker is weaker and more diffused compared to the signal from *pCLV3::H2B-mCherry*, and the *pTCSn::GFP* and *DR5rev::GFP* markers are cytoplasmic, rendering them unsuitable for spot-based segmentation. Therefore, the 3D models with these reporters were segmented into individual cells using FM4-64-stained membranes, and the mean intensity of the signal was calculated for each segmented cell. The decreasing size of the inflorescence meristem with developmental age (17, 19) poses a challenge for the comparative analysis of meristems at different ages. To overcome this issue, we focused on cells within the central zone of the meristem. The central zone of Arabidopsis shoot apical meristem has been defined as cells within a 30 µm perimeter from its center (35). To analyze comparable subsets of cells in the central zone, we focused on cells located within 21 µm radius of the meristem apex (Fig. 1g-j). We observed that this volume still excludes areas where emerging flower primordia start to protrude, even in the smallest arrested meristems. When comparing arrested meristems to those two weeks after bolting (2 wab), we observed that cells in arrested meristems are marginally smaller, resulting in a slightly higher cell count within the analyzed volume (Supplementary Fig. 3). However, this difference is negligible. Thus, the volume-based subset approach can be employed to facilitate the quantitative analysis of the same group of cells within each meristem, ensuring comparability of the data across different meristems.

### Age-dependent changes in inflorescence meristem

To assess the age-dependent changes in the inflorescence meristem, we quantified the expression of the stem cell marker *pCLV3::H2B-mCherry* in inflorescence meristems (IM) from 1 day after bolting (dab) until PA (Fig. 2a-e,u). Approximately 150-200 cells expressed *pCLV3::H2B-mCherry* in meristems from 1 dab to 2 wab, but the number decreased to less than 50 in the arrested meristem, consistent with its reduced size (Supplementary Fig. 4a). We also observed differences in the signal intensity between meristems of different ages. At 1 dab, *pCLV3::H2B-mCherry* expression was detected in a large area of the shoot apex with less variation in signal intensity between detected nuclei (Fig. 2a). This is the time when the inflorescence bolt is growing rapidly, but flowers have not yet opened. At 1 wab when the first flowers began to open, *pCLV3::H2B-mCherry* expression was strongest in the middle of the central zone and gradually decreased toward the peripheral zone (Fig. 2b). The signal intensity in the central zone further increased and reached a maximum at 2 wab (Fig. 2c). At PA, the *pCLV3::H2B-mCherry* was restricted to a small area of the central zone and signal intensity slightly decreased to levels comparable to meristems at 1 wab (Fig. 2d,u). This indicates that the stem cell niche is maintained at PA, although its size is significantly reduced.

**Fig. 2.**
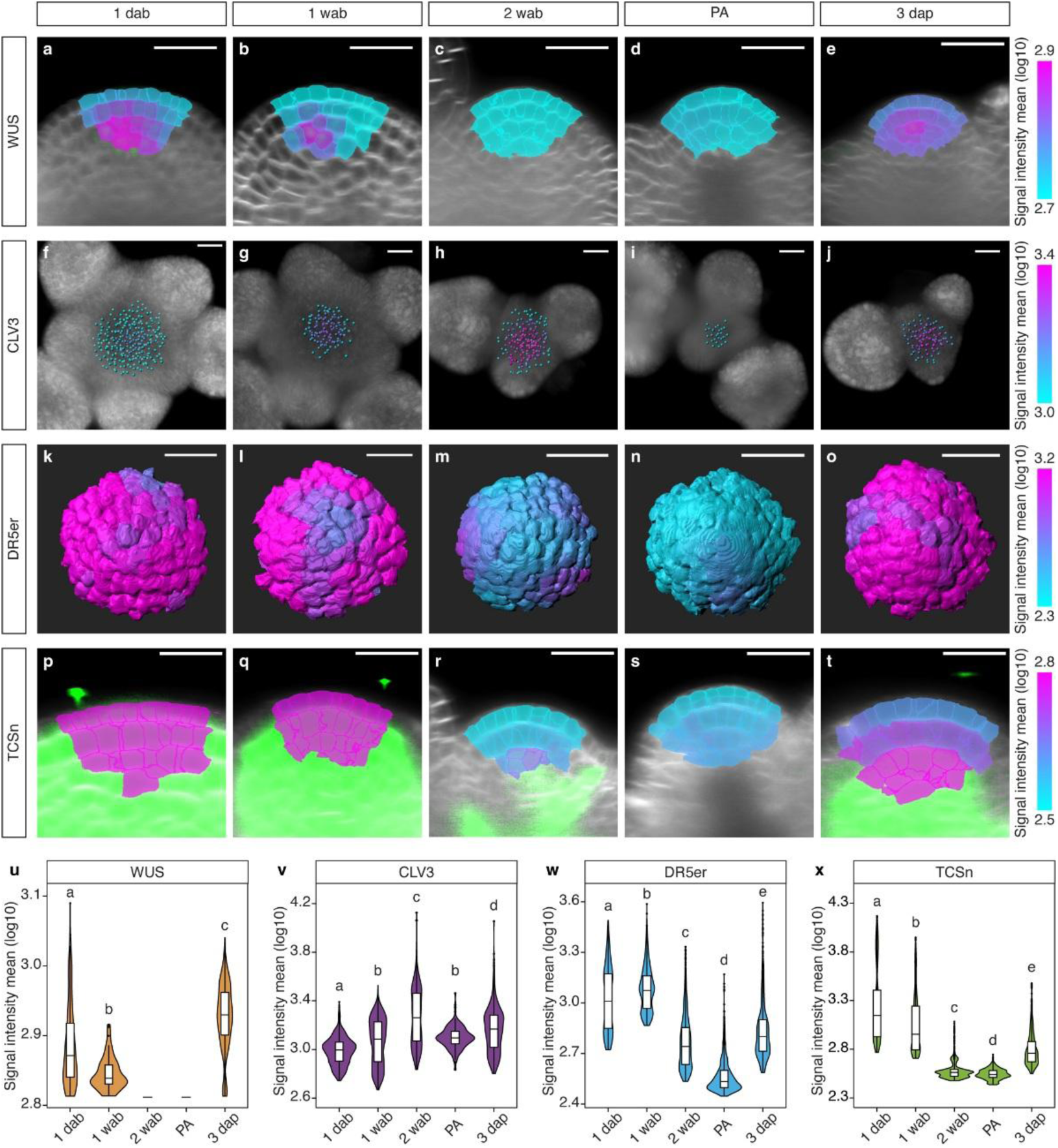
Age-dependent changes in inflorescence meristems. **(a-e)** Top views of 3D IM reconstructions with segmented *pCLV3::H2B-mCherry* nuclei. Mean intensity of the signal is shown by colormap. Plasma membrane is highlighted by *SYP132:GFP* (gray). **(f-j)** Longitudinal sections through IM models; colormap indicates mean intensity of the *pWUS::WUS-linker-GFP* signal in cells within 21 µm form the meristem apex. Plasma membrane is stained with FM4-64. **(k-o)** Longitudinal sections through IM models; colormap indicates mean intensity of *TCSn::GFP* signal in cells within 21 µm form the meristem apex. Green indicates *TCSn::GFP* signal outside the analyzed volume. Plasma membrane is stained with FM4-64. **(p-t)** Top views of segmented cells in 3D IM models that are within 21 µm form the meristem apex. Heatmaps indicate mean intensity of the *DR5rev::GFP* signal in each cell. Scale bars = 20 µm. **(u-x)** Quantification of mean signal intensity of *pCLV3::H2B-mCherry* (u), *pWUS::WUS-linker-GFP* (v), *TCSn::GFP* (w) and *DR5rev::GFP* (x) in IMs at 1 dab, 1 wab, 2 wab, PA and 3 days after pruning (dap) is shown by hybrid violin-box plots. In (v), a cutoff of 2.8 was applied to exclude the background signal. Charts with different lowercase letters are significantly different from each other (ANOVA, Tukey’s HSD test; p-value < 0.01, 215-1040 cells from 5 different meristems were included for each category).

We next analyzed the stem cell maintenance factor WUS using the *pWUS::WUS-linker-GFP* reporter. Expression of the WUS reporter was restricted to the deeper layers of the IM with the strongest signal at 1 dab, then its level decreased at 1 wab, and became undetectable at 2 wab (Fig. 2f-j,v). A similar pattern was also observed with the cytokinin signaling reporter *pTCSn::GFP*, which was more strongly expressed in the deeper cell layers compared to the uppermost layers (Supplementary Fig. 4d-f). The overall *pTCSn::GFP* signal, highest at 1 dap, gradually declined and became nearly undetectable by 2 wab; no signal was detected at PA (Fig. 2k-o,w). These data suggest that WUS expression and cytokinin signaling do not decrease just before PA, but start to decline already from early stages of inflorescence bolt formation.

Since inflorescence arrest was proposed to be caused by auxin (14, 23), we analyzed auxin signaling using *DR5rev::GFP* sensor. Reporter expression was primarily restricted to the L1 layer, with local maxima marking the initiation of floral primordia (Fig. 2p-t,x). Auxin signaling was high at 1 dab, its level peaked at 1 wab and then gradually declined reaching minima at PA. Nevertheless, even during PA, the *DR5rev::GFP* signal was detectable at the putative initiation sites of floral primordia (Supplementary Fig. 4b,c). These data show that PA is accompanied by a decrease in auxin signaling, which is preceded by a dampening of cytokinin signaling. These observations are consistent with the idea that emerging fruits gradually divert cytokinin transport from the IM, eventually rendering it susceptible to PA.

### Fruit removal rapidly re-establishes cytokinin signaling in arrested meristem

Removal of fruits reactivates the arrested meristem and restores cytokinin signaling and WUS expression (Fig. 2j,o,v,w). Removal of competing sink organs may restore the acropetal flow of cytokinin into the IM leading to its activation. Such an abrupt redirection of the cytokinin flow should result in a rapid resumption of cytokinin signaling in the IM. Indeed, we observed statistically significant increase in the *pTCSn::GFP* signal as early as two hours after pruning (hap), and the signal was very prominent at 6 hap (Fig. 3a-e). In roots, the *pTCSn::GFP* reporter senses cytokinin with a delay of 30-60 min after exogenous application of benzyl-aminopurine (Supplementary Fig. 5), suggesting that the level of cytokinin in the IM starts to increase 1-1.5 h after the fruit removal.

**Fig. 3.**
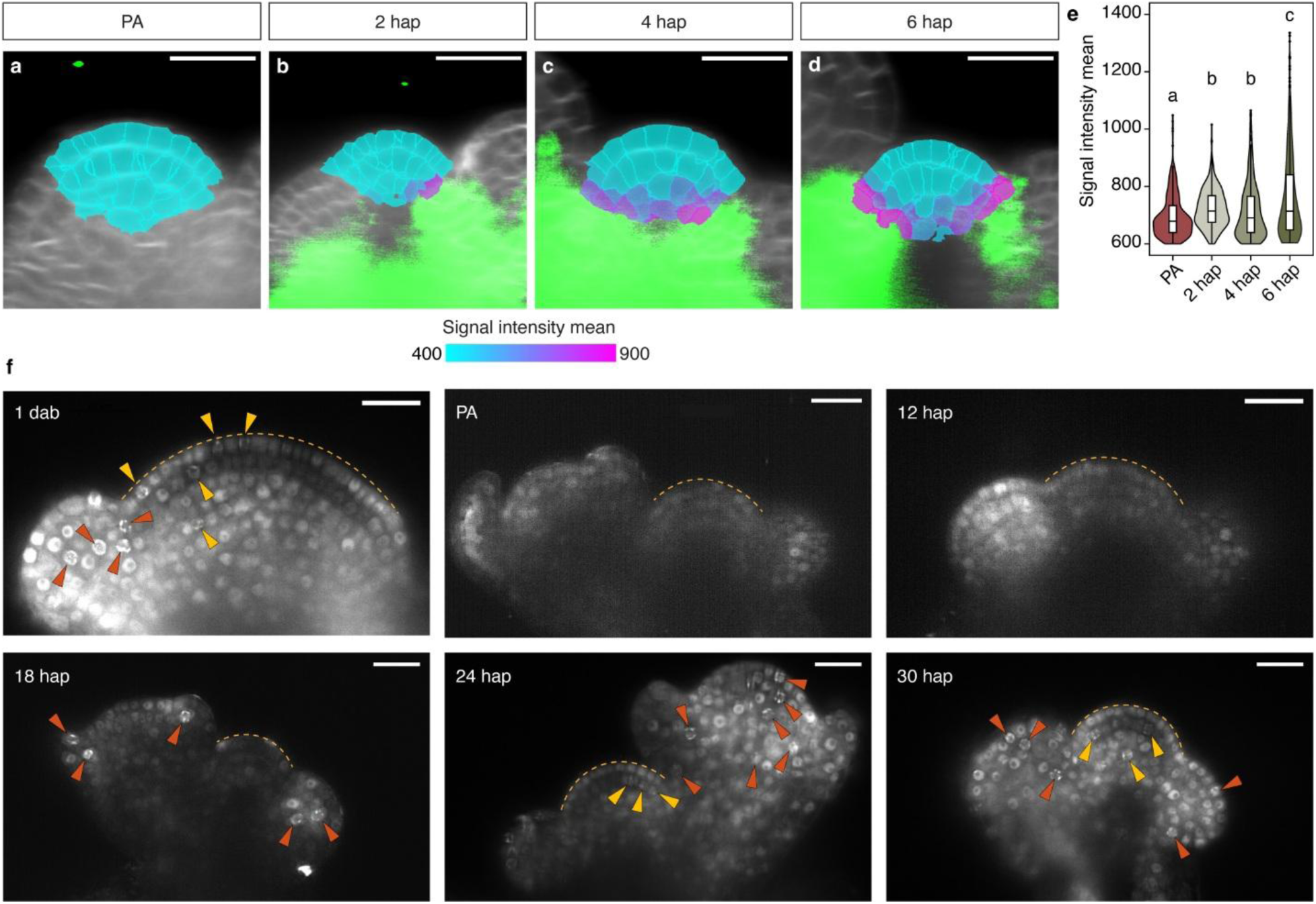
Reactivation of inflorescence meristem by pruning. **(a-d)** Longitudinal sections of IM reconstructions showing *TCSn::GFP* expression at PA (a), 2 hours after pruning (hap) (b), 4 hap (c) and 6 hap (d). Segmented cells within 21 µm form the meristem apex were color-coded according to heatmaps, indicating signal intensity mean. Cell membranes (gray) were highlighted using FM4-64. Scale bars = 20 μm. **(e)** Quantification of mean signal intensity of *TCSn::GFP* in samples as in (a–d) shown by hybrid violin-box plots. To exclude the background signal, an intensity cutoff of 600 was applied. Charts with different lowercase letters are significantly different from each other (ANOVA, Tukey’s HSD test; p-value < 0.01, 857-917 cells from 5 different meristems were included for each category). **(f)** Micrographs showing longitudinal sections of IMs with *PCNA:TagRFP* expression (gray). Inflorescences were imaged at 1 dab, PA, 12 hap, 18 hap, 24 hap and 30 hap and shown micrographs represented representatives from at least five IMs for each stage. A yellow dashed line outlines the IM dome. Arrowheads point to nuclei in S-phase located in inflorescence meristem (yellow) and flower primordia from stage P2 (orange). Scale bars = 20 μm.

IM reactivation leads to the resumption of the cell cycle, which can be monitored by the entry of cells into S-phase. To examine the timing of this process, we visualized the *PCNA:TagRFP* reporter, which is expressed in nuclei of proliferating cells and forms prominent nuclear speckles during S-phase (36, 37). While numerous S-phase nuclei were readily detected in the IM 1 dab, no such nuclei were visible during PA (Fig. 3f). Furthermore, the overall PCNA signal was low during PA, indicating a quiescent state of the meristematic cells. First S-phase nuclei were detected 18 hap in floral buds and 24 hap in IM indicating that floral buds resume growth before the IM.

### Absence of fruits keeps inflorescence meristems physiologically younger

Fruit production is a key determinant of longevity in many annual plants (4–7). In Arabidopsis, continuous removal of fruits by pruning or infertility caused by a mutation substantially delay the onset of proliferation arrest, extending the longevity of the IM and increasing flower production (Fig. 4a,b)(6). To assess how the absence of seeds affects inflorescence meristem physiology, we analyzed CLV3 expression along with auxin and cytokinin signaling in continuously pruned plants 2 wab. While inflorescence meristems of both pruned and untreated plants showed a similar number of CLV3 positive stem cells, the signal intensity was slightly lower in pruned plants (Fig. 4c-e). In contrast, auxin and cytokinin signaling were significantly increased by pruning 2 wab (Fig. 4f-k), with signals comparable to untreated meristems 1 wab (Figure 2w,x). These data suggest that inflorescence meristems in seedless plants are in a physiologically younger state than the age-matched meristems in fertile plants.

**Fig. 4.**
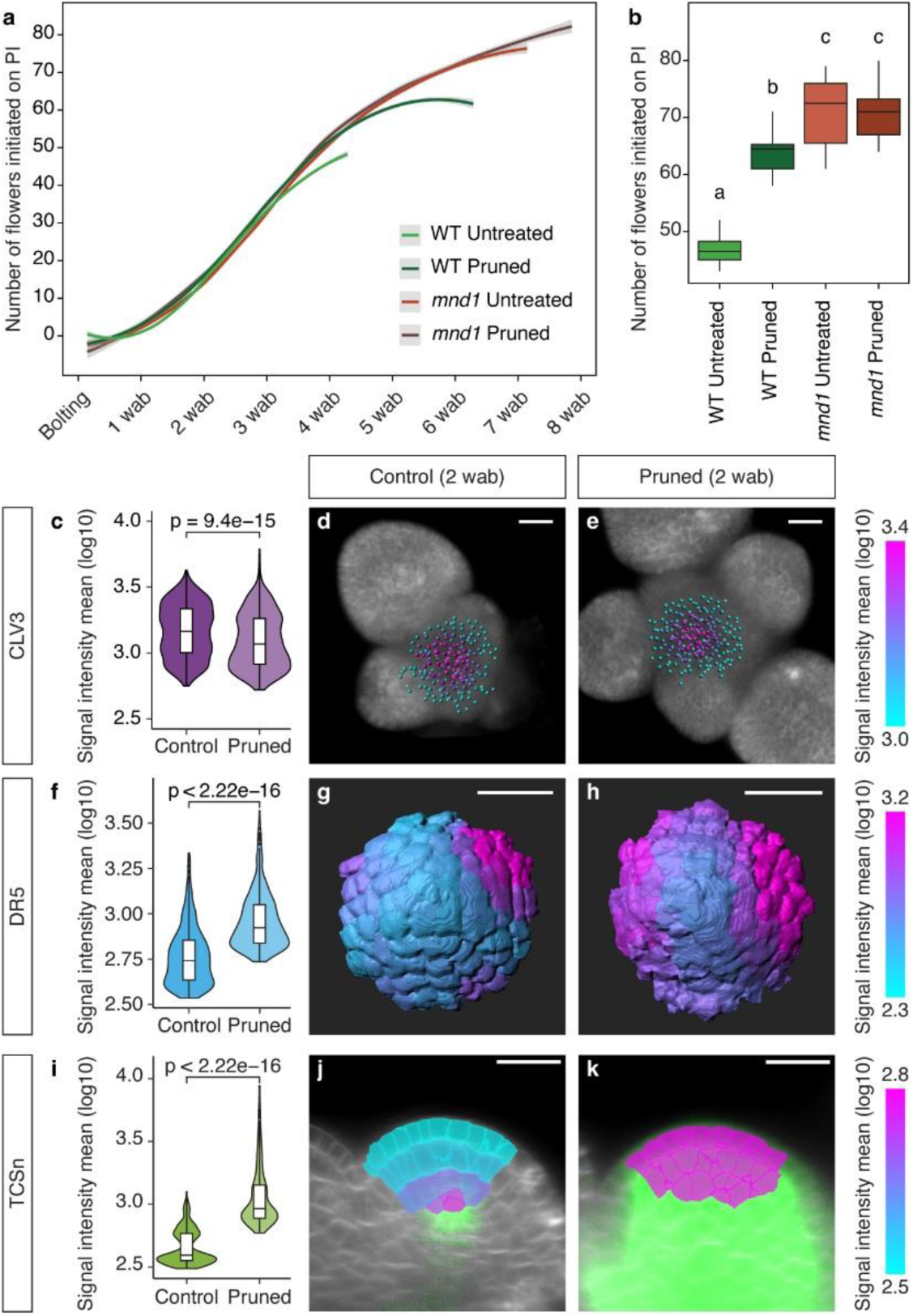
Effect of fruits on inflorescence meristem. **(a)** Trend lines showing the number of flowers initiated on PI in untreated and continuously pruned wild type and sterile *mnd1* mutants, respectively. Trend lines are calculated using the loess fitting method; grey areas represent confidence intervals of 0.95. n = 20 plants per treatment. **(b)** Box plot showing total number of flowers initiated on PI of samples as in (a). Box plots with different lowercase letters are significantly different from each other (ANOVA, Tukey HSD test; p-value < 0.01). **(c-e)** *pCLV3::H2B-mCherry* signal quantification in IM of pruned and untreated WT plants. (d,e) Top views of 3D IM reconstructions with segmented *pCLV3::H2B-mCherry* nuclei. Mean intensity of the signal is shown by heatmap. Plasma membrane is highlighted by *SYP132:GFP* (gray). **(f-h)** *DR5rev::GFP* signal quantification in IM of pruned and untreated WT plants. (g,h) Top views of segmented cells that are within 21 µm form the meristem apex. Heatmap indicates mean intensity of the *DR5rev::GFP* signal in each cell. **(i-k)** *TCSn::GFP* signal quantification in IM of pruned and untreated WT plants. Longitudinal sections through IM models; heatmap indicates mean intensity of *TCSn::GFP* signal in cells within 21 µm form the meristem apex. Green indicates *TCSn::GFP* signal outside the analyzed volume. Plasma membrane stained with FM4-64 is indicated in gray. Scale bars = 20 µm. Hybrid violin-box plots indicate quantification of signal intensity mean of *pCLV3::H2B-mCherry* (c), *DR5rev::GFP* (f) and *TCSn::GFP* (i) (714-919 cells from 5 different meristems were included for each category). The statistical significance of differences between pruned and untreated plants was calculated using the Wilcoxon signed-rank test.

### Manipulation of cytokinin levels in bolting plants affects the onset of PA

The quantitative imaging data (Figs. 2w; 3e, 4i) show a strong anticorrelation between seed formation and cytokinin signaling in the IM, indicating a causative role of cytokinin in linking reproductive effort to PA. This is supported by the observations that mutants with aberrant cytokinin signaling show pronounced changes in the onset of inflorescence arrest (15, 20, 25, 26). However, these mutants display pleotropic vegetative growth phenotypes, which complicates the interpretation of the observed effects on the flower production. To overcome this issue, we used an inducible system to modulate the cytokinin levels in adult plants. We took advantage of plants expressing either CYTOKININ OXIDASE 2 from barley, which deactivates cytokinin, (*proCaMV35S>GR>HvCKX2*), or bacterial ISOPENTENYL TRANSFERASE, which mediates its biosynthesis (*proCaMV35S>GR>ipt*), from the dexamethasone (DEX) inducible promoter (38). Transgene expression in these plants was induced by application of DEX solution to roots at different stages of bolting. Induction of *proCaMV35S>GR>ipt* resulted in prolonged flowering and differentiation of more flowers (Fig. 5a,b). In contrast, induction of *proCaMV35S>GR>HvCKX2* accelerated the inflorescence arrest and reduced flower formation, with the earlier DEX application having the greater effect (Fig. 5c,d).

**Fig. 5.**
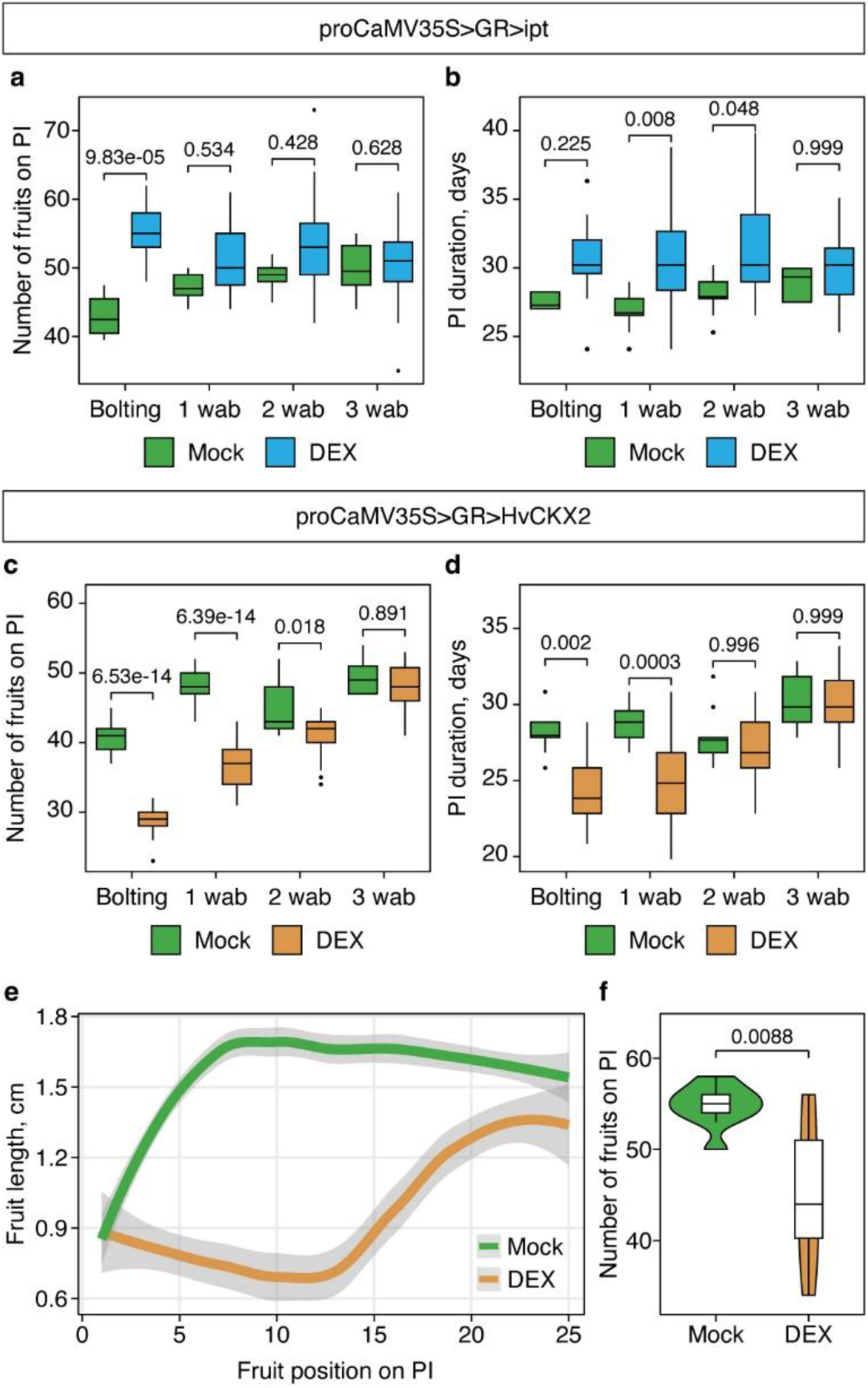
Effect of cytokinin levels on PA. **(a,b)** Box plots showing effect of *proCaMV35S>GR>ipt* induction on number of fruits on PI (a) and PI duration (b) after treating plants at bolting, 1, 2 and 3 wab either with mock (n = 9, 9, 9 and 8 plants, respectively), or with DEX (n = 15, 23, 23 and 22 plants, respectively). **(c,d)** Box plots showing effect of *proCaMV35S>GR>HvCKX2* induction on number of fruits on PI (a) and PI duration (b) after treating plants at bolting, 1, 2 or 3 wab either with mock (n = 10, 9, 9 and 10 plants, respectively), or with DEX (n = 17, 21, 21 and 22 plants, respectively). **(e)** Trend lines showing the length of fruits formed on PI as indicative of fertility of proCaMV35S>GR>HvCKX2 plants after local treatment with mock (n=11 plants) or DEX (n=10 plants). Trend lines are calculated using the loess fitting method; grey areas represent confidence intervals of 0.95. **(f)** Violin-box plots showing effect of proCaMV35S>GR>HvCKX2 induction on number of fruits on PI in the same plants as in (e). Statistical significance of differences between the group means in (a-d, f) was calculated using the Wilcoxon signed-rank test.

To further test whether local cytokinin deprivation in the IM impacts the PA, we sprayed the apex of the main inflorescence bolt of *proCaMV35S>GR>HvCKX2* plants with DEX at 1 day after bolting and monitored flower and fruit production until PA. The local DEX treatment abolished seed formation in subsequently formed siliques (Fig. 5e), supporting the notion that cytokinin is required for placental activity and ovule development (24). Fertility on the main inflorescence bolt was restored only in later-developed flowers (Fig. 5e). Notably, despite the reduced seed set, PA occurred earlier in DEX treated plants compared to mock-treated controls (Fig. 5f). This observation indicates that local cytokinin deprivation in shoot apex overrides the effect of infertility on PA.

### Role of cytokinin transport in fruit development and inflorescence arrest

This experiment strongly supports the idea that cytokinin represents a systemic signal that regulates the activity of IM. Cytokinins are primarily produced in roots and distributed via xylem to aboveground tissues (30, 31). To test whether cytokinin can be transported shootwardly into the IM, we applied trans-Zeatin (tZ) in lanolin to the internode between 20^th^ and 21^st^ fruit on the primary bolt during PA. This led to the induction of *TCSn::GFP* signal in arrested IM and surrounding floral buds 48 hours after the treatment (Fig. 6a-d).

**Fig. 6.**
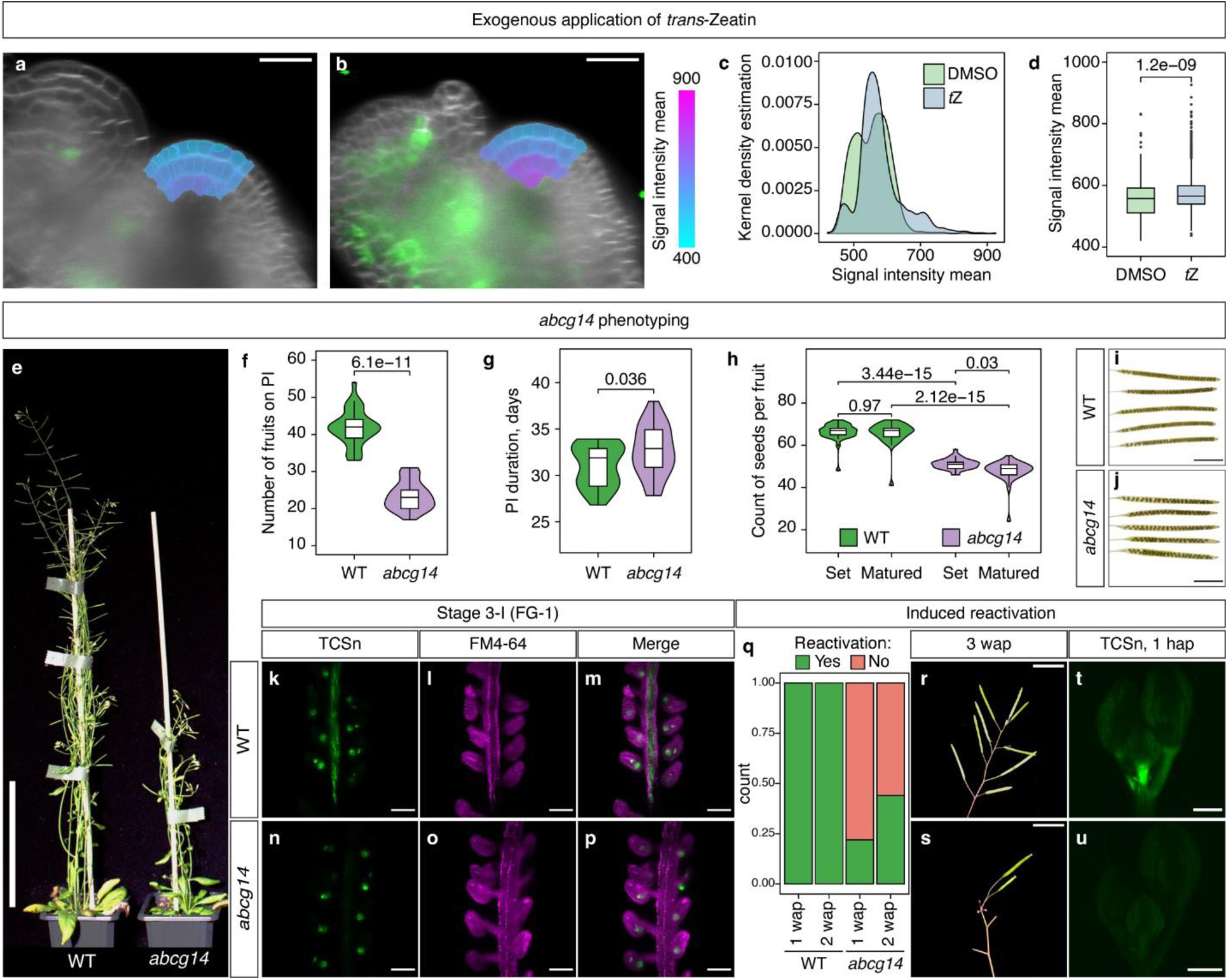
Effect of cytokinin transport on PA. **(a,b)** Longitudinal sections of IMs expressing *TCSn::GFP* (green) from inflorescences treated at PA by application of DMSO (a) or 300 μM *trans*-Zeatin (b) in lanolin paste. The lanoline paste was applied to the inflorescence stem internode separated by 20 fruits from unopened buds and meristems were imaged 48 hours after treatment. Heatmap indicates mean intensity of the *TCSn::GFP* signal in cells within 21 µm form the meristem apex. Plasma membrane stained with FM4-64 is indicated in gray. Scale bars = 20 µm. **(c,d)** Quantification of mean signal intensity of *TCSn::GFP* in IMs of *t*Z treated inflorescences shown in kernel density (c) and box plots (d). The statistical significance of differences between the mean of groups was calculated using the Wilcoxon signed-rank test (n = 828 and 858 cells combined from five different meristems for DMSO and *t*Z treated samples, respectively). **(e)** Representative picture of 45-days-old wild type (Col-4) and *abcg14* plants. Scale bar = 10 cm. **(f,g)** Violin-box plots showing the number of fruits on PI (f) and PI duration (g) in wild type and *abcg14*. The statistical significance of differences between the mean of groups was calculated using the Wilcoxon signed-rank test (n = 29 plants for each genotype). **(h)** Violin-box plot showing set of ovules and matured seeds per fruit in wild type and *abcg14*. The statistical significance of differences between mean of groups was calculated using the Wilcoxon signed-rank test (n = 30 fruits at position 5-10 on PI from total from 5 different PIs). **(i,j)** Images of cleared mature fruits from the same position on PI of wild type (m) and *abcg14* (n). Scale bar = 5 mm. **(k-p)** Micrographs showing MIPs of pistils from wild type and *abcg14* plants expressing *TCSn::GFP* (green). Pistils were imaged with ovules at stage FG-1 (k–m) and with embryos at the pre-globular stage (n–p). Cell membranes (magenta) were highlighted using FM4-64. Scale bar = 100 μm. **(q-s)** Reactivation of IM in *abcg14* plants. (q) Bar charts showing fraction count of induced reactivation of PIs in wild type and *abcg14* plants (n = 50 plants per group). (r,s) Representative pictures showing reactivated meristems in wild type (r) and *abcg14* mutant (s). Scale bar = 5 mm. (t,u) Micrographs showing MIPs of inflorescences at 1 hour after pruning from wild type and *abcg14* plants expressing *TCSn::GFP* (green). Scale bar = 300 um.

We next asked whether cytokinin is also transported into emerging fruits. Developing seeds are among the tissues with the highest cytokinin levels, and cytokinins are key regulators of the seed size and number (32, 39, 40). While it is well established that cytokinin biosynthesis occurs in developing seeds, the contribution of transport from maternal tissues has been less explored (32). To address this question, we analyzed plants deficient in cytokinin long-distance transport mediated by ATP-binding cassette transporter subfamily G14 (ABCG14) (41, 42). Arabidopsis *abcg14* mutants have severely impaired root-to-shoot translocation of tZ-type cytokinins and exhibit aberrant growth (41, 42) (Fig. 6e). We also found that the number of fruits on PI was significantly reduced in *abcg14* compared to wild type, while the duration of flowering was unaffected (Fig. 6f,g). This can be attributed to an increased plastochron ratio, as also reported for mutants with impaired cytokinin production (43). In addition, consistent with the role of cytokinin in regulating placental activity (24), *abcg14* mutants showed a reduced number of ovules and mature seeds (Fig. 6h-j). Examination of cytokinin signalling using *TCSn::GFP* showed a very strong signal in developing ovules and embryos in both wild type and *abcg14* plants (Fig. 6k-p), reaffirming that local cytokinin biosynthesis occurs from the earliest stages of female reproductive development. Nevertheless, *TCSn::GFP* was also observed in the replum and placenta during ovule development in wild type, but not in *abcg14* mutants (Fig. 6k-p). These results suggest that ABCG14 mediates cytokinin transport into fruits and that this process contributes to the seed formation.

We next analysed the reactivation of arrested IM in *abcg14* mutants. While fruit removal readily induced the formation of fertile flowers in the wild type, less than 50% of *abcg14* mutants showed any signs of IM reactivation 2 wap (Fig. 6q-s). Furthermore, the reactivated *abcg14* meristems rarely develop over 2 fertile siliques. We also observed that *abcg14* mutants exhibit substantially reduced *TCSn::GFP* expression in the inflorescence apex (Fig 6t,u) showing that ABCG14 is critical for cytokinin transport into shoot apex. These data support the idea that reactivation of arrested IMs following fruit removal is mediated by the redirection of ABCG14-meidated cytokinin transport to IM.

### Nitrate availability modulates the onset of inflorescence arrest

Flower formation and subsequent reproductive effort must be carefully coordinated with mineral nutrient availability of to ensure successful reproduction and resource allocation. Cytokinins act as signaling molecules that coordinate growth and developmental responses to nitrate availability (43, 44). To test whether nitrate levels affect proliferative arrest and flower production, we grew plants on sand and watered them with nutrient solution containing 5 mM KNO3 as the sole source of nitrogen. After bolting, plants were treated with nutrient solution containing different concentrations of KNO3 (0, 2.5, 5, 7,5 or 10 mM) until PA (Fig. 7a). We observed a dose-dependent response to nitrate concentration in the duration of flower production (Fig. 7b,c). We also observed a similar response in the number of rosette inflorescences that were initiated after bolting (Fig. 7d). Unlike primary and rosette inflorescences, the number cauline inflorescences (CI) was not affected (Fig. 7e) suggesting that nitrate-starved plants preferentially release axillary buds on already developed inflorescences, rather than in the rosette.

**Fig. 7.**
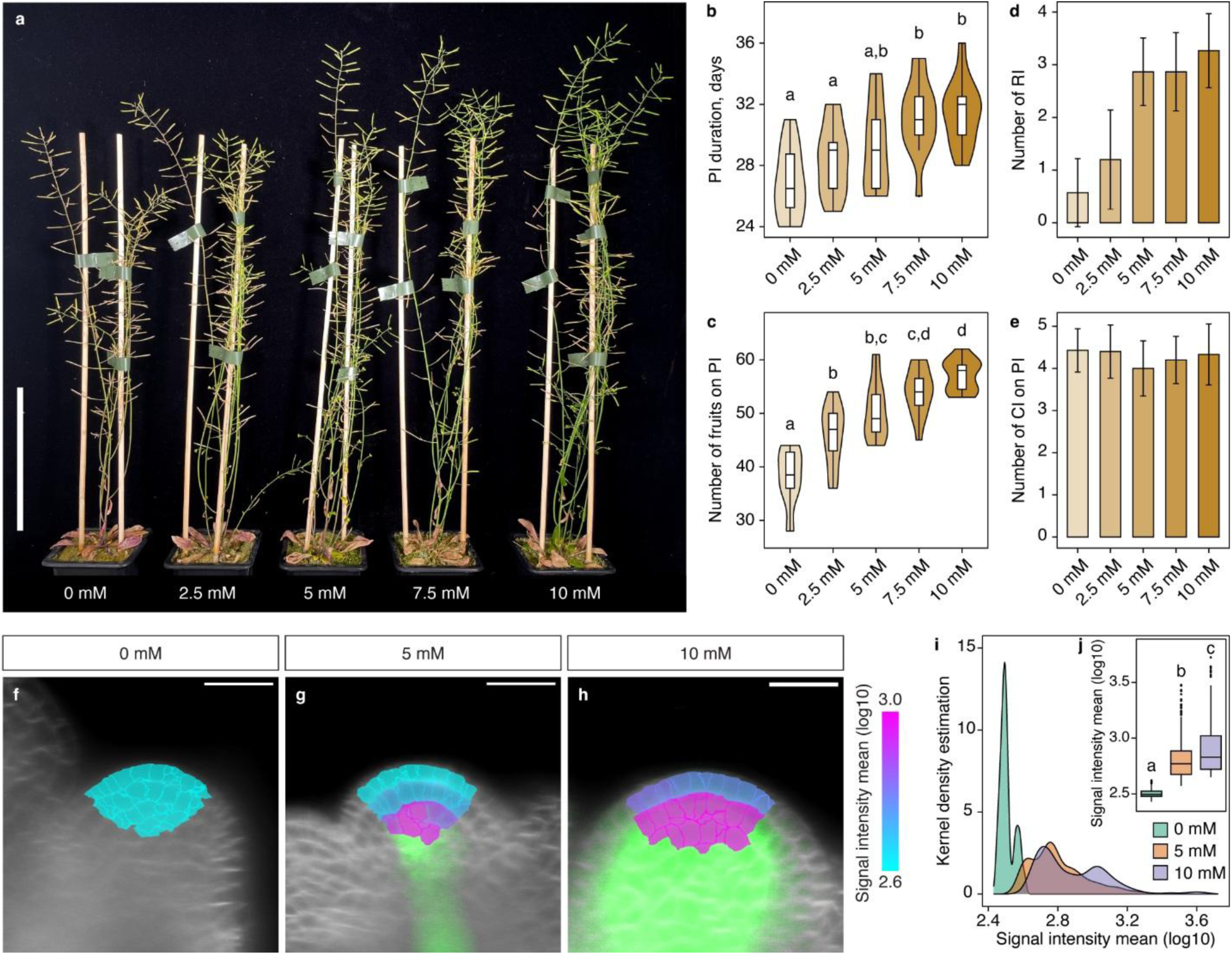
Effect of nitrate levels on meristem arrest. **(a)** A representative picture of plants grown on substrate supplemented with different KNO3 concentrations. The picture was taken when plants of all treatment groups reached PA. Scale bar = 10 cm. (**b,c)** Violin-box plots showing PI duration (b) and number of fruits on PI (c) in plants grown on different KNO3 concentrations. Plots with different lowercase letters are significantly different from each other (ANOVA, Tukey HSD test; p-value < 0.01, n = 15 plants per treatment). **(d,e)** Bar charts showing mean number of rosette inflorescences (RI) (d) and cauline inflorescences (CI) on PI (e) in plants of treatment groups as in (a). Error bars represent standard deviation (n = 15 plants per treatment). **(f-h)** Longitudinal sections of IMs expressing *TCSn::GFP* (green) from plants grown on different concentrations of KNO3 and imaged 2 wab. Heatmap indicates mean intensity of the *TCSn::GFP* signal in cells within 21 µm form the meristem apex. Plasma membrane stained with FM4-64 is indicated in gray. Scale bars = 20 µm. **(i-j)** Quantification of the mean signal intensity of *TCSn::GFP* in IMs from plats grown at different concentrations of KNO3 shown in kernel density and box plots (inset). Plots with different lowercase letters are significantly different from each other (ANOVA, Tukey HSD test; p-value < 0.01; n = 877, 784 and 808 cells combined from five different meristems from plants grown at 0 mM, 5 mM and 10 mM KNO3, respectively).

To validate that these growth responses are related to changes in cytokinin signaling, we examined *TCSn::GFP* expression in inflorescence meristems of plants supplemented with nutrient solution containing 0, 5 or 10 mM KNO3 for two weeks after bolting (Fig. 7f-j). Plants deprived of nitrate (0 mM KNO3) showed no *TCSn::GFP* expression in the IM, whereas an increasing signal was observed in meristems watered with increasing concentrations of KNO3. These data suggest that the duration of reproductive growth is modulated by nitrate availability using root-derived cytokinin as a long-distance messenger.

## Discussion

Reproduction is a resource-demanding process that in plants is supported by massive relocation of nutrients from vegetative tissues, often leading to leaf senescence and demise of an entire organism (45, 46). Therefore, the ability to precisely coordinate the timing and duration of reproduction with developmental and environmental cues is central to the evolutionary success of a species. In some plants, the duration of flowering is determined by the activity of the inflorescence meristem, which is subject to feedback control from seeds already produced. Such regulation must integrate information about ongoing fruit and seed production, as well as nutrient availability and overall plant physiology. Recent studies in Arabidopsis suggest that plant reproductive effort is monitored by auxin flux from newly emerging fruits (14, 23). This process reduces auxin canalization from the adjacent developing floral buds, inducing their growth arrest and dormancy.

Several studies have indicated a role of cytokinin in regulating inflorescence arrest and flower production (15, 19, 20, 24–27). Here we provide evidence that cytokinin functions as a systemic signal that determines the physiological status and proliferative capacity of the inflorescence meristem by integrating information on reproductive effort and nutrient availability. We suggest that in line with the “nutrient drain” hypothesis, the partitioning of shootwardly transported cytokinin into developing fruits reduces its flux to the inflorescence meristem and ultimately induces the cessation of its activity.

Several lines of evidence support this scenario. First, our data from *abcg14* mutants indicate that developing fruits act as a sink for cytokinin transport (Fig. 6k-p). Second, cytokinin signalling in the IM gradually declines as new fruits are formed (Fig. 2), and this process is delayed by continuous fruit removal (Fig. 4). Third, reactivation of an arrested meristem by pruning is accompanied by a rapid activation of cytokinin signalling, possibly due to the redirection of cytokinin transport caused by the removal of competing sinks (Fig. 3). This is further substantiated by the observation that cytokinin transport mutants are impaired in meristem reactivation (Fig. 6q-u). Finally, manipulation of cytokinin levels through inducible deactivation or biosynthesis during bolting has a pronounced effect on the onset of meristem arrest (Fig. 5). Importantly, local cytokinin deactivation mitigates the effect of reduced fertility on proliferation arrest, providing a strong support to the idea that cytokinin flow represents the mechanism linking plant reproductive status with meristem activity.

The timing of inflorescence arrest and flower production are also affected by other physiological parameters, such as carbohydrate status, photo-thermal conditions and nitrate concentration (Fig. 7) (14, 47). Cytokinin regulates growth responses to nitrate (43, 44) and thus may also integrate inputs on nutrient availability to fine-tune plant reproductive effort.

The correlative inhibition of plant growth by fruits and seeds is a well-documented phenomenon with significant practical implications, observed in numerous plants (11). Two alternative scenarios, known as the “death hormone” and “nutrient drain” hypotheses, have been proposed to explain these observations. These scenarios are not necessarily mutually exclusive. Auxin may function as a “death hormone” by being canalized basipetally from developing seeds, providing information about fruit production near the inflorescence meristem. Conversely, shootwardly-mobile cytokinin may serve as a more systemic signal that is “drained” by newly emerging sink organs, thus providing a broader perspective on plant physiological status and reproductive effort. Together, these mechanisms suggest a complex interplay of hormonal signaling that coordinates growth and reproduction in plants, allowing adaptive responses to changing environmental and developmental conditions.

## Methods

### Plant material and growth conditions

All plants used in the study were *Arabidopsis thaliana* ecotype Col-0, with the exception of the *abcg14* mutant which is in the Col-4 background. Plants were grown in a growth chamber at 21⁰C with 50-60% humidity under 16h/8h light/dark cycles (light intensity 150 µmol m^-2^ s^-1^) in a 3:1 by volume mixture of soil:vermiculite. The following Arabidopsis lines were used in the study: *pCLV3::H2B-mCherry* (48), *pWUS::WUS-linker-GFP* (49), *DR5rev::GFP* (50), *TCSn::GFP* (51), *PCNA:TagRFP* (36), *proCaMV35S>GR>HvCKX2* (52), *proCaMV35S>GR>ipt* (38), and *abcg14* (41). In the original *pWUS::WUS-linker-GFP* line, the construct complemented vegetative defect of *wus* mutant (GABI-Kat line GK870H12) phenotype, but plants were semi-sterile. Because sterility and semi-sterility affects flower production, we outcrossed the *wus* mutation and the resulting line in wild type background, which did not show any growth or fertility defects, was analyzed.

### Plant treatments

The *proCaMV35S>GR>HvCKX2* and *proCaMV35S>GR>ipt* plants were induced at the indicated time points by watering each plant with 50 ml of 10 µM dexamethasone. Control plants were watered with 50 mL of 0.0001 % (v/v) DMSO. For local treatment, plants were induced by direct application of 10 µl of 10 µM dexamethasone to inflorescence at 1 dab; control plants were treated with 0.0001 % (v/v) DMSO. For cytokinin application to the inflorescence bolt, trans-zeatin (10 mM in DMSO) was mixed with lanolin paste at a ratio of 3:100 (v/v) and the paste was applied to the internode between fruits 20 and 21 counted from the top cluster of arrested buds. For the nitrate resupply experiment, seedlings were germinated on soil for 7 days, transferred to individual pots with the sand/zeolite mixture in the ratio 3:2 (v/v) pre-watered with 50 mL of nutrient solution (53) containing 5 mM KNO3, 2 mM MgSO4, 3 mM CaCl2, 2.5 mM KH2PO4 (adjusted to pH 5.5), 70 µM H3BO3, 14 µM MnCl2, 0.5 µM CuSO4, 1 µM ZnSO4, 0.2 µM NaMoO4, 10 µM NaCl, 0.01 µM CoCl2, 50 µM Fe-EDTA. Plants were fed with 25 mL of the nutrient solution weekly until bolting. From bolting, plants were watered weekly with 50 mL of the nutrient solution supplemented with either 5 mM KCl (0 mM NO3), or 2.5 mM KNO3, 2.5 mM KCl (2.5 mM NO3), or 5 mM KNO3 (5 mM NO3), or 7.5 mM KNO3 (7.5 mM NO3), or 10 mM KNO3 (10 mM NO3).

### Phenotyping of reproductive growth

We used nomenclature as in Ware et al. (23): primary inflorescence is produced by the primary embryonic shoot apex; secondary inflorescences are formed from axillary buds in the axils of cauline leaves (cauline inflorescence) and rosette leaves (rosette inflorescence). For the developmental timing data plants were assessed daily for beginning of bolting (scored upon appearance of visually detectible primary inflorescence from leaves rosette) and inflorescence arrest (scored when there are no more open flowers on inflorescence). The number of fruits and inflorescences was recorded after PA unless stated otherwise. Reproductive growth of inflorescence was defined as the period from the beginning of bolting to PA.

### Inflorescence meristem imaging

The inflorescence meristem was dissected with sharp tweezers under a stereomicroscope in a drop of ½ MS medium supplemented with sucrose (5%); all pedicels with flower buds were carefully removed. For plasma membrane staining, a drop of fresh FM^TM^4-64 dye (Invitrogen, ref. number T13320; concentration 0.1 mg/mL) was applied to a glass slide and the tissue was stained for 2 min. Without prior washing, the IM was inserted into a glass capillary (Brand: size 4, blue, ref. number 701910) filled with cooled but liquid mounting medium (½ MS 5 % sucrose with 1% low melting agarose) containing well vortexed fluorescent beads (1 µl of beads per 200 µl of the media; Invitrogen, FluoSpheres^TM^ polysterene, size 1 µm; red (580/605) ref. number F13083; blue/green (430/465) ref. number F13080), and the medium was solidified for 5 min. Microscopy was performed with a Light-sheet Z.1 microscope (Zeiss). The capillary was inserted into the microscope chamber containing liquid 5xMS medium (36) and the sample was slightly pushed out of the capillary into liquid medium of the chamber. Imaging was performed using 20x objectives (Detection optics 20x/1.0), dual side illumination (illumination optics 10x/0.2) and two-track imaging with 488 nm laser for GFP/YFP, and 561 nm laser for TagRFP/mCherry/FM4-64.

### Image processing

Raw data in .czi format were deconvolved in ZEN (Zeiss) using the regularized inverse filter, strength 2. During this step, images form dual side illumination were fused to a single image. Deconvolved .czi files were imported into FIJI and using the Multiview reconstruction plugin (54) Z stacks from all 8 angles were registered and aligned using the fluorescent bead signal (Fast descriptor-based, translation invariant model to regularize with affine). The same registration was then applied to the second channel, which did not contain the fluorescent bead signal from. All angles were then fused together and exported in .ics/.ids format. Using the Imaris file converter (ImarisFileConverter 9.2.1), the fused image was converted to .ims format and opened in the Imaris software (Imaris 9.2.1). Segmentation was performed based on signal localization. For the cytoplasmic signal, the segmentation algorithm “cells” was selected and using the wizard, membranes were selected for segmentation (cell smallest diameter 2.5 µm, automatic cell membrane threshold). The surface was generated from segmented cells. A reference frame was then manually positioned at the apex and a filter for 21µm distance from the reference frame was then applied to the surfaces. Signal artifacts present outside of the meristem were manually curated. The pixel intensity mean from the channel containing the cytoplasmic signal was then exported from the clean filtered selection. For the *pCLV3::H2B-mCherry* nuclear signal, the segmentation algorithm “spots” was selected and 2 µm spheres were detected directly in the analysed channel. Due to the low experimental background, all spots with the threshold 0 were included and therefore no filtering was necessary, as we were confident that they all belonged to meristematic tissue. Values of pixel intensity mean from all spots were exported. Fluorescence intensity of *TCSn::GFP* expression in roots was measured as described (55).

### Data analysis

For each meristem, values of the mean signal intensity for each segmented cell/spot (hereafter “data point”) and the distance of the data point from a reference frame in x-y-z were exported (Supplemental Tables 1-5), and analyzed by statistical tests specified in the figure legends using R. For each age group or treatment condition, 5 inflorescence meristems were imaged and their data points were pooled for analysis (Supplemental Table 6). Signal intensity was plotted in R using ggplot2 (https://doi.org/10.1007/978-3-319-24277-4_9). 3D scatter plots representing models of individual IMs analyzed in this study are provided in the Supplemental Data Set. These 3D plots were generated using Plotly (https://github.com/plotly/plotly.R).

## Supporting information

Supplemental Dataser

## Acknowledgements

This work was funded by the Czech Science Foundation (23-07969X) and by the Ministry of Education, Youth, and Sports of the Czech Republic from the project TowArds Next GENeration Crops, reg. no. CZ.02.01.01/00/22_008/0004581 of the ERDF Programme Johannes Amos Comenius. We acknowledge the support from CEITEC MU Core facilities Plant Sciences and CELLIM, supported by the Czech-Bioimaging (No. LM2018129) infrastructure project funded by MEYS CZ. We also thank Dr. Ortrun Mittelsten Scheid, Dr. Jan Lohmann and Dr. Helene Robert Boisivon for providing reporter lines used in this study. Finally, we are indebted to Dr. Tom Bennett and Dr. Vivek Raxwal for their valuable comments and discussion on this paper.

---

**Movie 1. 3D reconstruction of the Arabidopsis inflorescence meristem expressing *pCLV3::H2B-mCherry* 2 wab**. Nuclei expressing *pCLV3::H2B-mCherry* are indicated in red, the plasma membrane is visualized with *SYP132:GFP* (green).

**Movie 2. 3D reconstruction of the Arabidopsis inflorescence meristem expressing *TCSn::GFP* 1 dab**. The *TCSn::GFP* is indicated in red, the plasma membrane is visualized by FM4-64 staining (gray). The animation also shows segmented cells that were within 21 µm of the meristem apex.

