## Supplementary Figures 1-5 for "Cytokinin acts as a systemic signal coordinating reproductive effort in Arabidopsis"

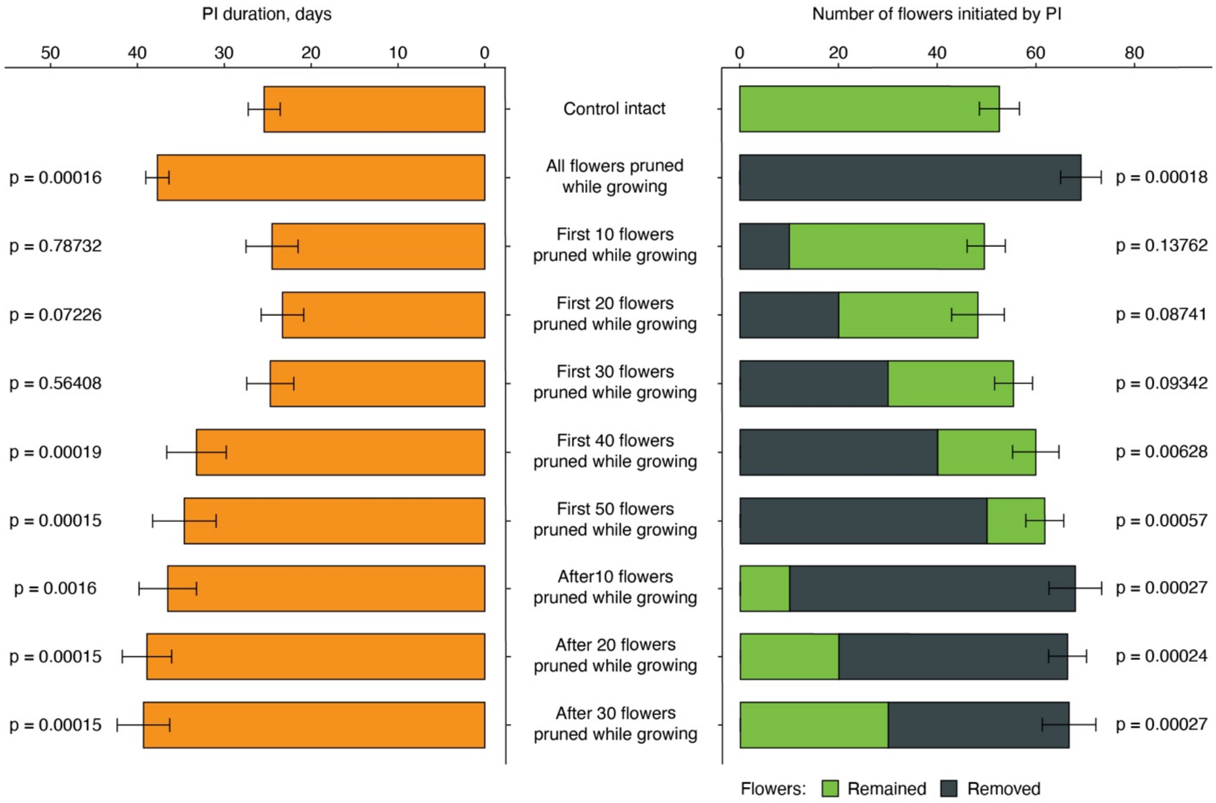


**Supplementary Fig. 1. Effect of pruning on inflorescence meristem arrest.** Bar charts showing mean primary inflorescence (PI) duration and the number of fruits on PI in plants pruned as indicated in the chart. Error bars represent standard deviation (n = 10 plants per group). The statistical significance of differences between the control and treatment groups was calculated using the Wilcoxon signed-rank test.

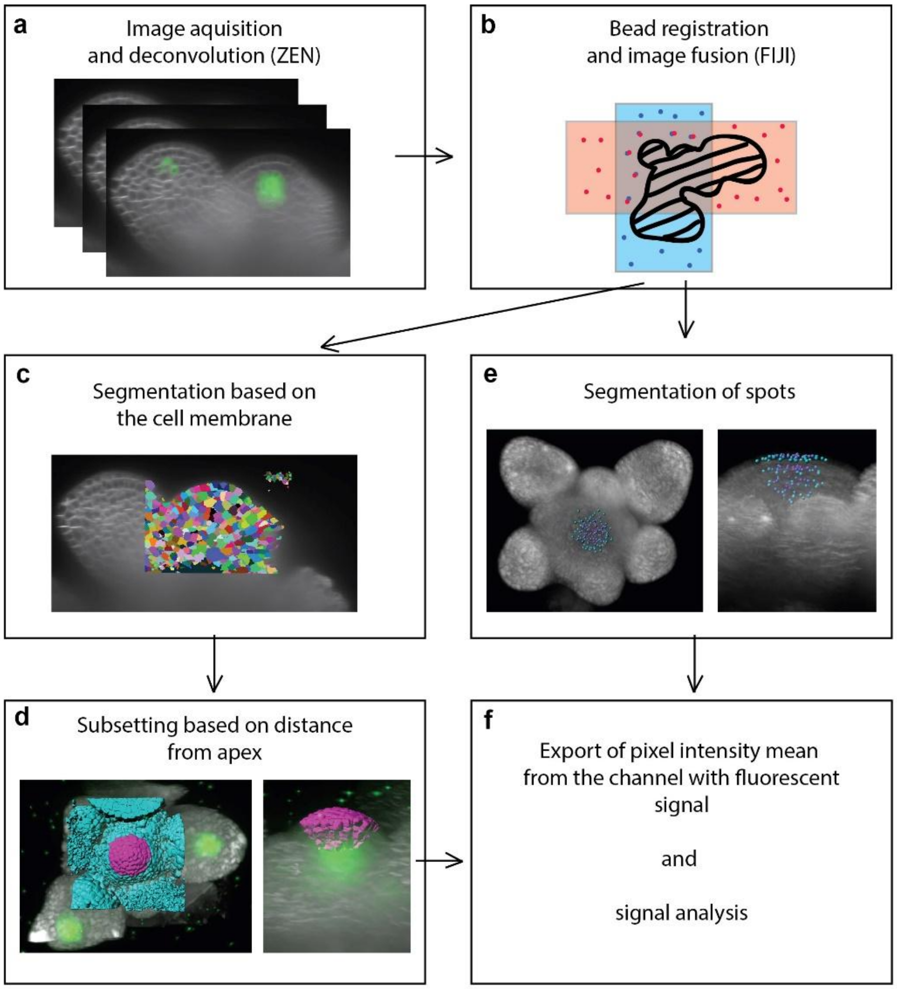


**Supplementary Fig. 2. Image processing pipeline. (a)** Raw data containing 16 Z-stacks, acquired from 8 different angles, were deconvolved using ZEN. **(b)** Deconvolved Z-stacks were registered and aligned to each other using fluorescent beads in FIJI and fused into a single Z-stack containing signal from the entire sample. **(c)** For the cytoplasmic signal, cells were segmented based on the plasma membrane signal obtained by FM4-64 staining in Imaris and **(d)** cells located at 21 µm distance from the apex were selected for further analysis. **(e)** For the nuclear signal segmentation, we used the spots algorithm in Imaris and for each detected nucleus a sphere of 2 µm was included in the downstream analysis. **(f)** Pixel intensity mean values for all selected segments were exported and statistically analyzed.

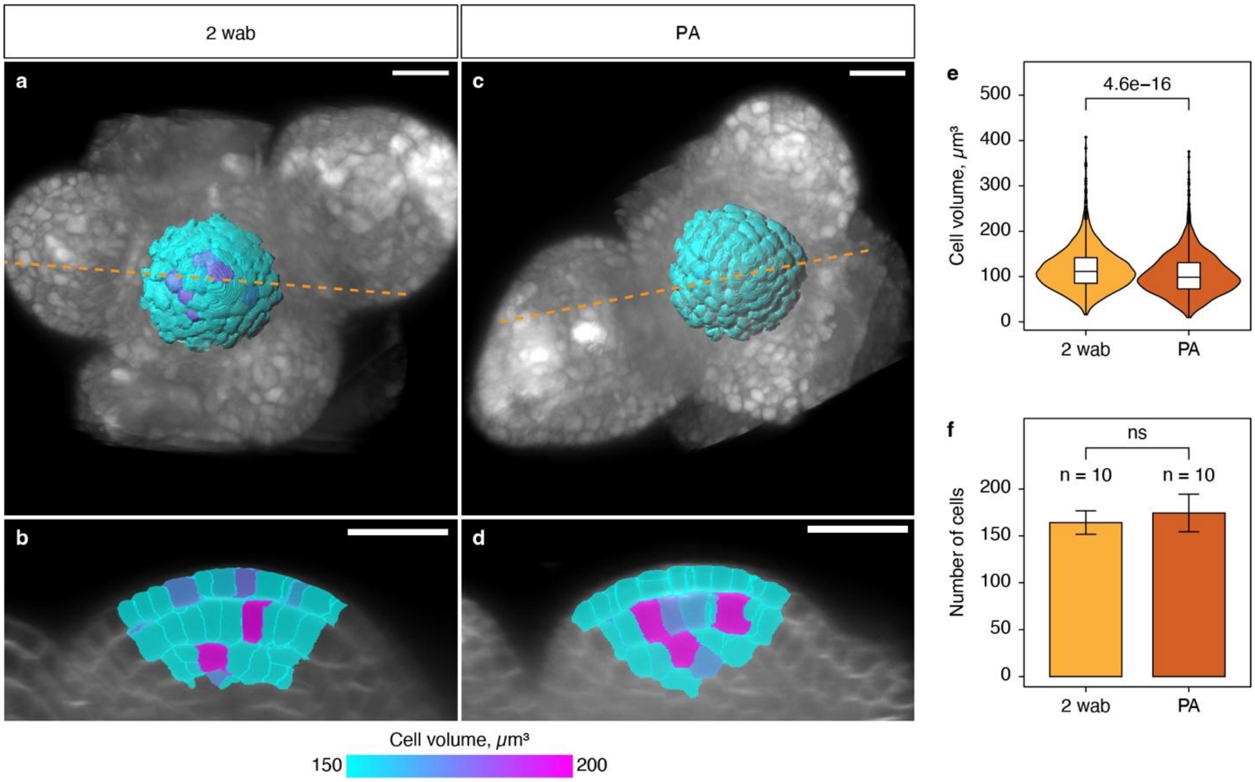


**Supplementary Fig. 3. Cell volume in the apex-proximal region of the inflorescence meristems. (a-d)** Top views (a,c) and longitudinal sections of 3D reconstituted IMs dissected at 2 wab and PA. Segmented cells within 21 μm of the meristem apex are highlighted with a heatmap indicating cell volume ($\mu m^3$). Cell membranes (gray) were highlighted using FM4-64. Scale bar = 20 μm. **(e)** Violin-box plot showing quantification of the cell volume ($\mu m^3$) within 21 μm of the meristem apex in IMs dissected at 2 wab or PA (n = 10). Statistical significance of differences between the group means was calculated using the Wilcoxon signed-rank test. **(f)** Bar graph showing the mean number of cells within 21 μm of the meristem apex in IMs dissected at 2 wab or PA. Error bars represent standard deviation (n = 10). Statistical significance of differences between the group means was calculated using the Wilcoxon signed-rank test.

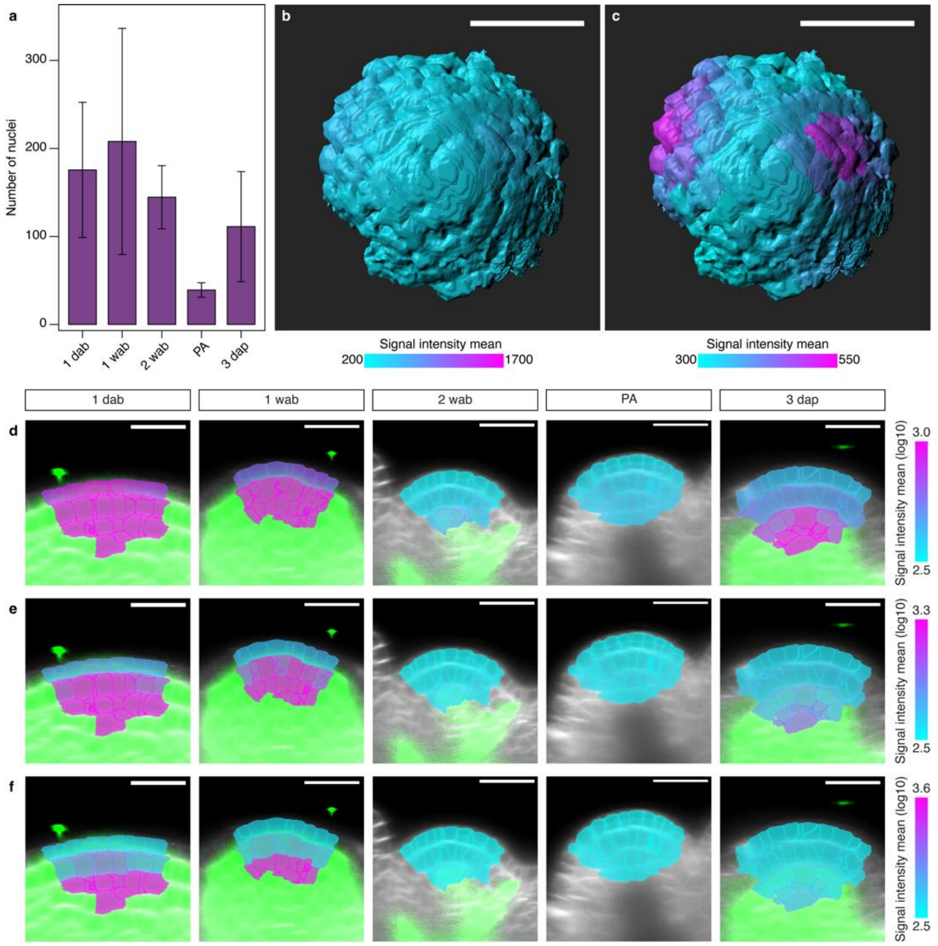


**Supplementary Fig. 4. CLV3 expression and auxin signaling in IMs at proliferative arrest. meristems. (a)** Bar graph showing the mean number of nuclei expressing *pCLV3::H2B-mCherry* in IMs of a different ages. Error bars represent standard deviation (n = 5 IMs). **(b,c)** Top views of 3D reconstructed and segmented IM with color rendering indicating *DR5rev::GFP* expression (only regions within 21 μm of the meristem apex are shown). The segmented cells were color-coded using a heatmap reflecting the mean signal intensity (log10-transformed) of *DR5rev::GFP*. Different range of the signal intensity was used in the heatmap to highlight weakly expressing cells in (c). Scale bar = 20 μm. **(d-f)** Longitudinal sections through the same sets of IM models with colormaps spanning different ranges of mean intensity of *TCSn::GFP* signal in cells within 21 μm from the meristem apex. The different scales of colormaps were used to highlight cytokinin signal pattern in this region. Green indicates *TCSn::GFP* signal outside the analyzed volume. Plasma membrane is stained with FM4-64. Scale bar = 20 μm.

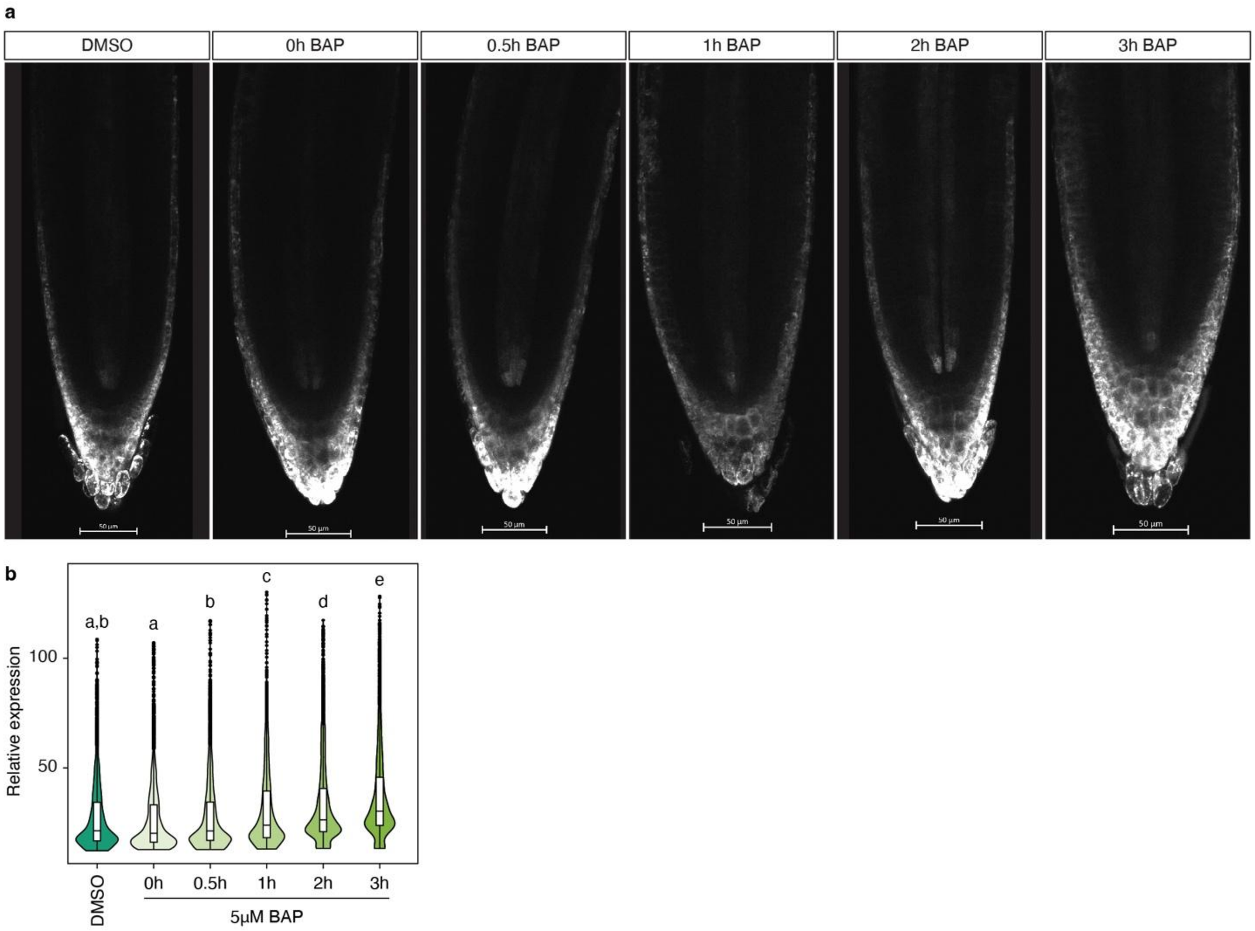


**Supplementary Fig. 5. Response time between cytokinin application and TCSn::GFP detection in roots. a**, Micrographs showing primary root tips of *TCSn::GFP* plants at different time points after treatment with 5 μM benzylaminopurine (BAP). Scale bar = 50 μm. **b**, Violin-box plot showing quantification of *TCSn::GFP* expression. Violon-boxes with different lowercase letters are significantly different from each other (ANOVA, Tukey's HSD test; p-value < 0.01; n = 10 roots per treatment).
